# D27-LIKE1 carotenoid isomerase has a preference towards *trans*/*cis* and *cis*/*cis* conversions in Arabidopsis

**DOI:** 10.1101/2022.04.06.487258

**Authors:** Zsolt Gulyás, Blanka Moncsek, Kamirán Áron Hamow, Pál Stráner, Eszter Badics, Norbert Incze, Éva Darkó, Valéria Nagy, András Perczel, László Kovács, Vilmos Soós

## Abstract

Carotenoids are colourful isoprenoids that contribute to a variety of physiological processes in plants. They also function as biosynthesis precursors of abscisic acid (ABA) and strigolactones (SLs). SL biosynthesis starts with the enzymatic conversion of all-*trans*-β-carotene to 9-*cis*-β-carotene by the DWARF27 (D27) isomerase. In Arabidopsis, D27 has two closely related paralogs, D27-LIKE1 and D27-LIKE2 which were predicted to be β-carotene-isomerases. Here we characterised D27-LIKE1 and identified some key aspects of its function. Arabidopsis *d27-like1-1* mutant does not display any SL or karrikin-deficient traits, however, it exhibits a substantially higher 9-*cis*-violaxanthin content. *In vitro* feeding assays with recombinant D27-LIKE1 revealed that the protein exhibits affinity to all β-carotene isoforms but with an exclusive preference towards *trans/cis* conversions and the interconversion between 9-cis, 13-cis and 15-*cis*-β-carotene forms. Feeding experiments with zeaxanthin and violaxanthin isomers revealed that D27-LIKE1 accepts these xanthophylls as substrates. The remarkably higher 9-*cis*-violaxanthin content of the mutant is accompanied by a slightly higher ABA level. Finally, we presented evidence that *D27-LIKE1* mRNA is phloem mobile and D27-LIKE1 is an ancient isomerase with long evolutionary history. In summary, we demonstrated that D27-LIKE1 is a carotenoid isomerase with multi-substrate specificity and has a characteristic preference towards the catalysation of *cis/cis* interconversion of carotenoids. Therefore, D27-LIKE1 is a potential regulator of carotenoid *cis* pools and eventually, SL and ABA biosynthesis pathways.

## Introduction

Generally, carotenoids are well known as pigments responsible for the myriads of colours of flowers, fruits and vegetables. Plant carotenoids are tetraterpenoid derivates, and classified into two groups based on their chemical structure (Ruiz-Sola and Rodríguez-Concepción, 2012). Carotenes are hydrocarbons while xanthophylls contain oxygen heteroatoms in different functional groups (Maoka, 2020). Carotenoids occur in *cis* and *trans* isomers in nature. The dominant isomers are the all-*trans* forms in plants, but extreme heat, high light or oxidative environment can induce spontaneous *trans/cis* isomerisation. For instance, β-carotene isomers occur in the following order: all-*trans*>9-*cis*>13-*cis*>15-*cis* (Guo, Tu and Hu, 2008). In addition to being pigments, carotenoids have many other essential roles in higher plants. They absorb the light in the blue-green region contributing to the increase of photosynthetic efficiency (Green, Anderson and Parson, 2003). However, this light-harvesting role of these compounds is dwarfed by their photoprotective ability (Green, Anderson and Parson, 2003). During high light conditions, carotenoids can prevent the photo-damage of photosynthetic apparatus through the operation of the violaxanthin-antheraxanthin-zeaxanthin cycle (VAZ cycle). When the photodamage of light harvesting complex is imminent, violaxanthin is converted reversibly to zeaxanthin via antheraxanthin by violaxanthin de-epoxidase (Müller, Li and Niyogi, 2001). Zeaxanthin promotes the thermal energy dissipation of excess absorbed energy through non-photochemical quenching (NPQ) (Müller, Li and Niyogi, 2001). Furthermore, carotenoids serve as precursors for apocarotenoid biosynthesis, which includes plant hormones such as abscisic acid (ABA) and strigolactones (SLs) (Nambara and Marion-Poll, 2005; Xie, Yoneyama and Yoneyama, 2010). ABA has essential role in several aspects of plant growth (seed development, dormancy, germination etc.), promotion of stomatal closure and various stress responses (Chen *et al*., 2020). ABA synthesis starts from the C_40_ precursor β-carotene (Nambara and Marion-Poll, 2005). In plastids, zeaxanthin-epoxidase converts zeaxanthin to all-*tran*s-violaxanthin (Audran *et al*., 2001). Then, *trans*-violaxanthin can either be converted to *trans*-neoxanthin by neoxanthin synthase (NXS) which is then converted to 9-*cis*-neoxanthin by a subtle isomerase or alternatively, the *trans*-violaxanthin can be converted to 9-*cis*-violaxanthin by an unknown mechanism (Finkelstein, 2013; Neuman *et al*., 2014). Nevertheless, both conversions require ABA4 and NXD1, although their proper function either as an isomerase enzyme or as a cofactor/auxiliary protein remained elusive (Neuman *et al*., 2014; Perreau *et al*., 2020). The 9-*cis* isomers of violaxanthin and neoxanthin are the subjects of oxidative cleavage by 9-*cis*-epoxicarotenoid dioxygenases (NCEDs) which produces C_15_ xanthoxin (Kalladan et al., 2019). Next, xanthoxin is converted to ABA by successive oxidative steps via abscisic aldehyde in the cytosol (Finkelstein, 2013).

Other important apocarotenoid-derived hormones are the SLs which are involved in controlling a wide range of plant developmental processes, including root architecture, mycorrhiza, stature and shoot branching, seedling growth and leaf morphology (Al-Babili and Bouwmeester, 2015; Waters *et al*., 2017). Enhanced shoot branching, lateral and adventitious root development and reduced stature are induced by SL deficiency (Stirnberg et al., 2002; Rasmussen et al., 2012). SLs are synthesised via a sequential oxidative cleavage of the precursor all-*trans*-β-carotene derivatives. First, DWARF27 (D27) catalyses the isomerisation of all-*trans*-β-carotene and the resulting 9-*cis-*β-carotene is cleaved by MAX3 and MAX4 dioxygenases (Alder *et al*., 2008). The SL precursor carlactone is then transported through the xylem and biologically active SLs are formed by MAX1 (Al-Babili and Bouwmeester, 2015) and LBO (Brewer et al., 2016).

The first committed step in SL biosynthesis is catalysed by the iron-containing D27 enzyme, which has been first identified in rice (Lin *et al*., 2009). *In vitro* studies with recombinant protein revealed that D27 mediates a reversible interconversion of all-*trans*-β-carotene towards its 9-*cis* isomeric form, which is the direct precursor of SL biogenesis (Alder *et al*., 2012). *D27* was shown to be induced by auxin, phosphate starvation and ABA, and lower levels of ABA have been observed in *d27* mutant rice and Arabidopsis (Abuauf *et al*., 2018; Haider *et al*., 2018; Liu *et al*., 2020), indicating a crosstalk with the ABA pathway. Having a common biosynthetic origin from carotenoids (Matusova *et al*., 2005), it is tempting to speculate that ABA and SL biosynthesis interfere with each other. It is still elusive whether D27 solely responsible for the synthesis of 9-*cis*-β-carotene (Waters et al., 2012) and, along with its paralogs, is involved in the isomerisation of ABA synthesis precursors (Perreau et al., 2020). Phylogenetically, the D27 protein family consists of three paralogs with a long history of evolutionary divergence (Waters, Brewer, *et al*., 2012). Two paralogs, At1g64680 (AtD27-LIKE1) and At4g01995 (AtD27-LIKE2) have no assigned function yet. In this study, we have investigated D27-LIKE1 in Arabidopsis, which was previously identified as a target of stress-related miRNAs in wheat (Cao *et al*., 2019). The high structural similarity enabled us to draw a parallel between the phylogenetic, physiological and enzymatic features of D27 and D27-LIKE1 and to provide ideas to further explore the function of D27-LIKE1.

## Results

### Ancestry of the DWARF27 family

The onset of large-scale sequencing projects enabled us to resolve the origin of DWARF27 family proteins with a higher accuracy (Leebens-Mack *et al*., 2019). First, protein sequences of D27-LIKE accessions from numerous algae and land plant species have been retrieved from ONEKP (www.onekp.com) and NCBI databases with the highest possible taxonomic coverage (Supplementary Table S1). The HMMER algorithm (Finn *et al*., 2015) predicted a conservative, 85 aa long domain (EMBL-European Bioinformatics Institute). These D27 domain sequences were used to infer the phylogenetic analysis using the NJ method (Fig 1.). Previous studies suggested that the DWARF27 family consists of three unambiguous and coherent clades, *D27-LIKE2, D27-LIKE1* and *D27*. Our basic tree topology partially agrees with these findings. A neo-functionalisation of an ancient algal *D27-LIKE* led to the emergence of the common ancestor of *D27-LIKE2* and *D27-LIKE1*. The monophyletic *D27-LIKE2* superclade separated early and present in all land plants but absent in algae suggesting that this superclade is a novelty in the green lineage. Interestingly, *D27-LIKE2* superclade splits into two distinct clades of which Clade Ia consists of all land plants excluding Angiosperms, which solely consists of Clade Ib. The duplication of the ancestral *D27-LIKE* gave rise to Superclade II containing the Angiosperm *D27-LIKE1* and *D27*. A series of duplication events led to a vast paralogy of *D27-like* sequences in algae, then Superclade II splitted into distinct, monophyletic clades. Clade IIa sequences are present in green algae and land plants except seed plants from which these ancient *D27-LIKE* genes (hereafter termed as *D27-LIKE3*) were eliminated early when Gymnosperms and Angiosperms emerged. Clade IIb sequences are present in *Coccomixaceae* and *Chlorellaceae*, in all land plants except *Anthocerotopsida* and contains Arabidopsis *D27-LIKE1*. Next, Clade IIc emerged which contains the canonical Arabidopsis *D27* sequence. It is noteworthy that *D27* is absent in all algal lineages, ferns and Gymnosperms and present only in mosses *sensu lato* and Angiosperms.

**Figure 1.**
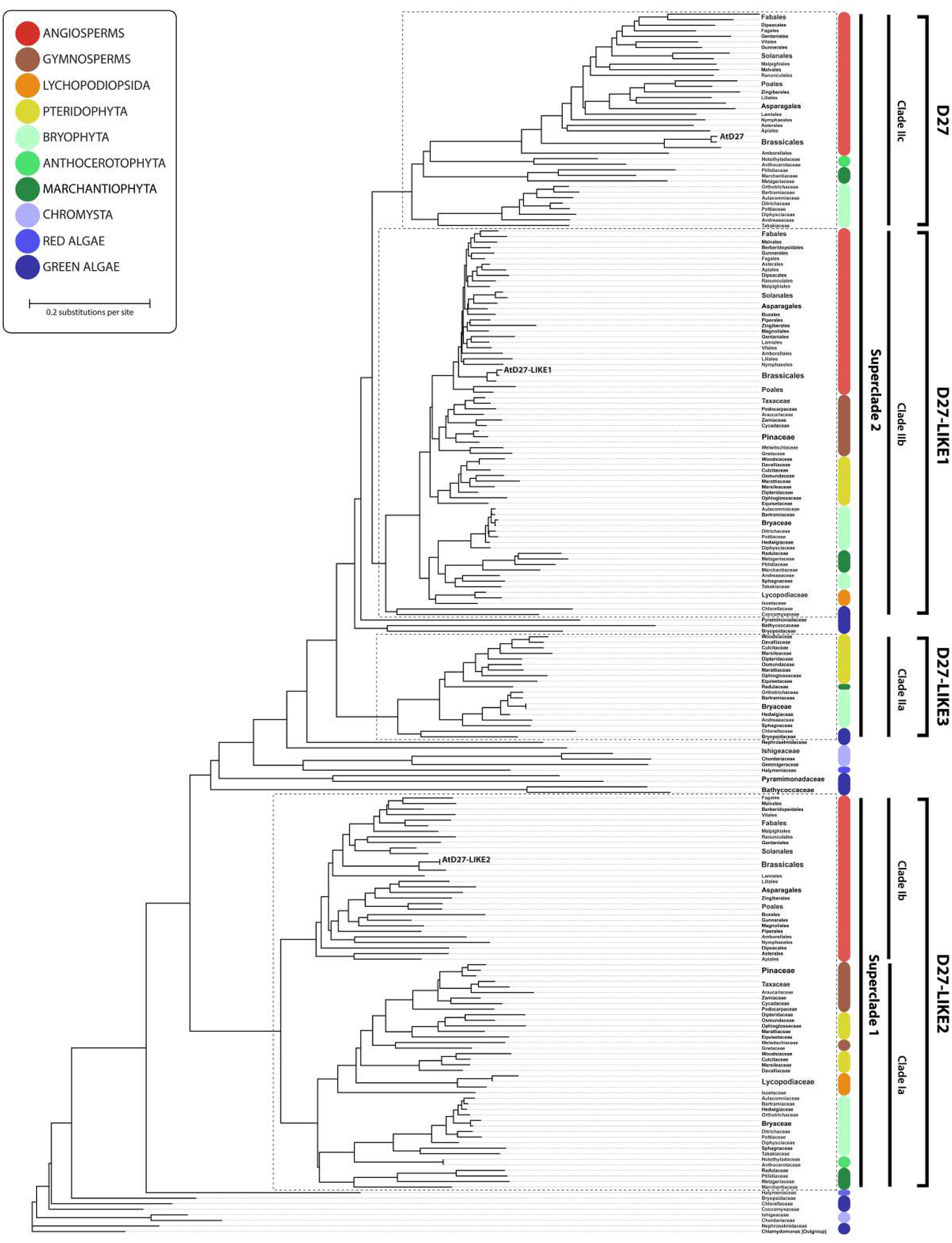
Reconstructed phylogenetic relationship of DWARF27 family proteins in the green plant lineage. HMMER-predicted D27-domain sequences (DUF4033) were extracted from the ONEKP (https://www.onekp.com) and NCBI (https://www.ncbi.nlm.nih.gov/) databases using Arabidopsis D27-like protein sequences as a query template. The tree was constructed using Geneious Prime 2021.2.2 by the neighbour-joining method (NJ), with bootstrap probabilities based on 1000 replicates. *Chlamydomonas reihardtii* D27-like protein sequence was used as outgroup. Arabidopsis D27 and its paralogs are indicated on the tree. The identifiers of each D27-like protein are listed in Supplemental Table S1.

The phylogenetic analysis enabled us to create a sequence logo to demonstrate conserved and specific aa residues in the four identified clades of *DWARF27* family in land plants (Supplementary Fig. S2). Interestingly, all four D27-LIKE domains are highly conservative. In sum, a higher resolution picture of D27 ancestry shows that the fourth member of the *DWARF27* family is absent in seed plants. Our tree topology suggests that *D27-LIKE1* and *D27* are sister branches and Gymnosperms unexpectedly lost *D27*.

### Arabidopsis D27-LIKE1 does not play a prominent role in SL or KAI2 ligand biosynthesis

To functionally characterise D27-LIKE1, we identified two available insertion mutant alleles (GT13552, GT22003) of *AtD27-LIKE1* (*D27-LIKE1*) in the L*er* genetrap collection (Martienssen, 1998). In GT22003, the insertion is present in the promoter of *D27-LIKE1*, and a reduced expression of *D27-LIKE1* has been detected, therefore the line has been excluded from the experiments. In GT13552, insertion is located within the first exon of *D27-LIKE1* (Fig. 2A) and the line displayed a severe lesion mimic phenotype, which segregated from the mutant allele when the plants were backcrossed to L*er*. After backcrossing three times, a line homozygous for GT13552 allele has been obtained and the progeny of this line has been termed as *d27-like1-1* (Fig. 2B).

**Figure 2.**
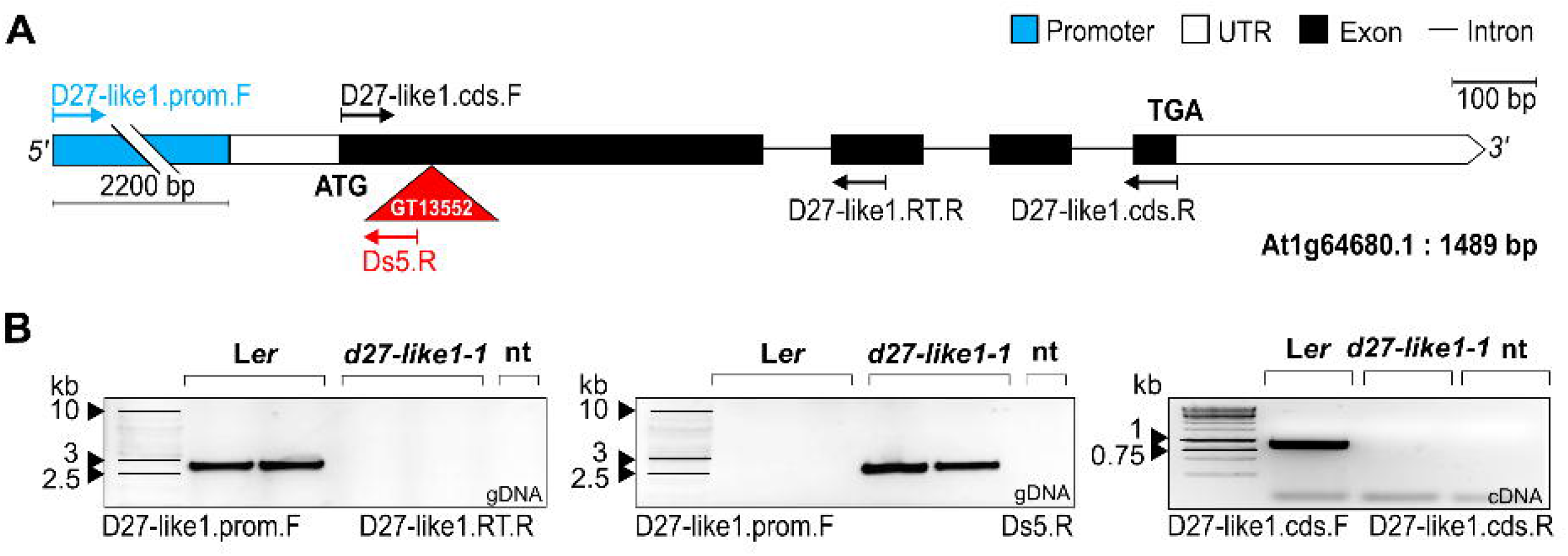
Gene model and genotyping of D27-L1KE1. A, Transcript structure of D27-LIKE1. The insertion in the GT13552 line is located in the first exon and is indicated by an inverted triangle. Arrows represent the orientation and position of primers, which were used for genotyping. UTR: Untranslated region. Bar: approximately 100 bp. B, RT -PCR analysis of backcrosscd *d27*-*like1-1* homozygous mutant plants (gDNA, cDNA). Primers used for genotyping are shown by arrows in “A”.

Next, the phenotypic traits related to the previously described paralogue *AtD27* (*D27*) were investigated. Arabidopsis *d27* mutants display increased branching phenotype with slightly lower plant height compared to the wt (Waters et al., 2012). Nonetheless, we did not observe such phenotypic traits in the *d27-like1-1* mutant (Fig. 3A, E and F). SLs have a significant role in leaf development suggested by previous observations in SL mutants (Bennett *et al*., 2016). We found no differences in leaf dimensions and leaf area between *d27-like1-1* and L*er* plants (Fig. 3B, G) suggesting that SL signalling is not perturbed in *d27-like1-1* plants. Next, we investigated whether karrikin signalling through KAI2, a paralog of D14, is affected in *d27-like1-1* plants. *rac*-GR24 and karrikin 2 (KAR2) supplementation in agar plates resulted in shorter hypocotyls of both wt and *d27-like1-1* seedlings (Fig. 3C, H). Karrikins are known to regulate root hair density and development via KAI2-mediated signalling pathway. We did not find a difference in root hair density between wt and *d27-like1-1* plants (Fig. 3D, I), suggesting that - together with the hypocotyl tests - *d27-like1-1* plants have no compromised karrikin signalling. These results indicate that D27-LIKE1 is not a prominent enzyme in SL or the elusive KAI2 ligand biosynthesis.

**Figure 3.**
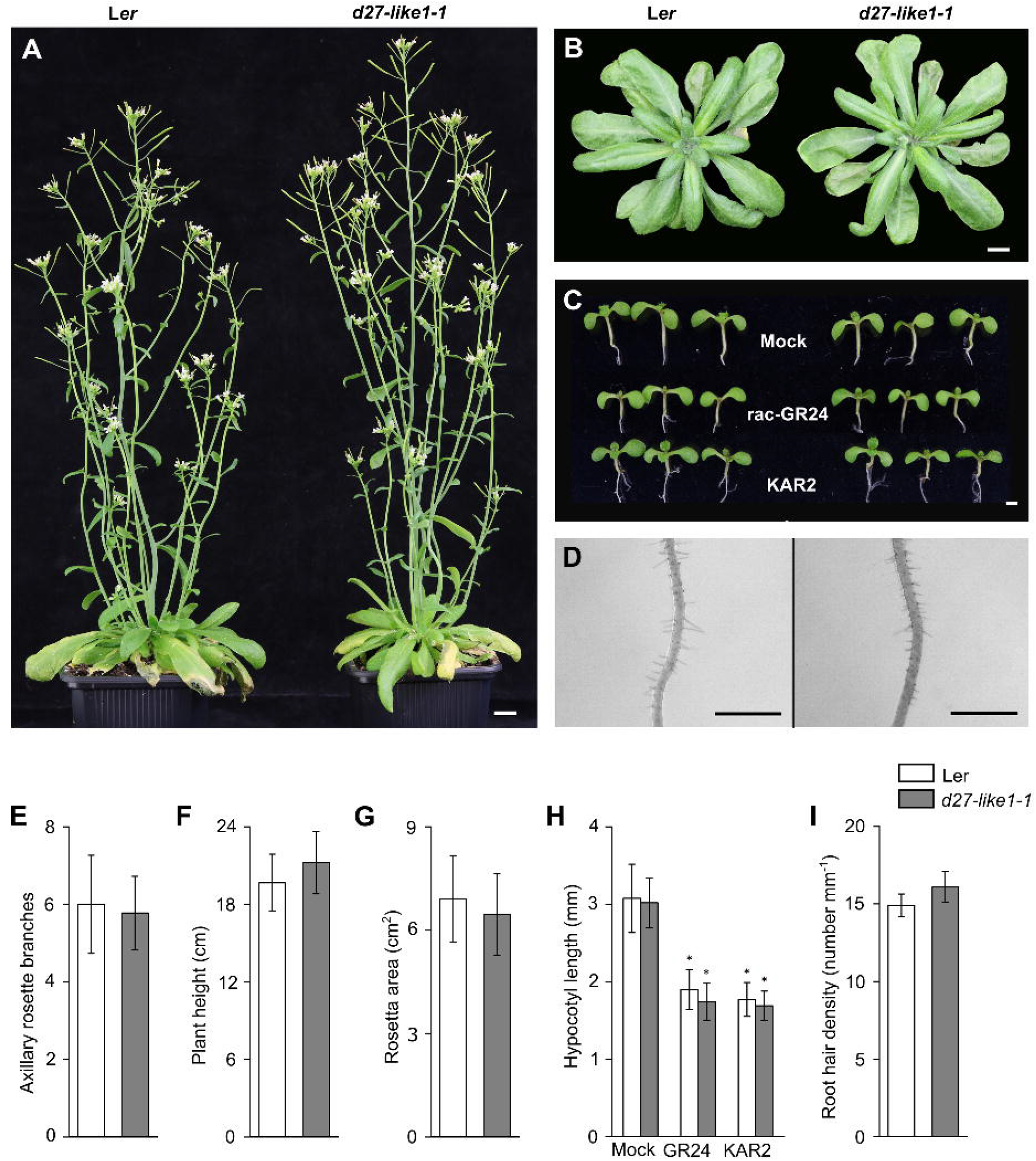
D27-LIKE1 does not have a prominent role in SL or KA12 ligand biosynthesis in Arabidopsis. A, Phenotype of wild -type L*er* (left) and *d27-like1-1* (right) adult plants. Plants shown are 6 -week-old and were grown under short day (8 h/16 h) conditions for 4 weeks and then under long day (16 h/8 h) conditions for 2 weeks Bar = 1 cm. B, 5-wcck-old rosettes of wild-type L*er* (left) and *d27-like1-1* (right) plants. Plants were grown under short day (8 h/16 h) conditions. Bar = 1 cm. C, Hypocotyl elongation of 10-day-old seedlings (L*er* left, *d27-like1-1* right) after 1 µM *rac*-GR24 and 1 µM KAR2 treatments. Bar = 1 mm. D, Root hair density of 10 -day-old seedlings (L*er* left, *d27-like1-1* right). Bars = 1 mm. E, Axillary rosette branches in 6-week-old L*er* and *d27-like1-1* plants. F, Plant height of 6-week-old L*er* and *d27-like1-1* plants described in A. G, Rosette area of 4 -week-old L*er* and *d27-like1-1* rosettes. Plants were grown under short day (8 h/16 h) conditions. H, Hypocotyl elongation of 7-day-old seedlings after GR24 and KAR2 treatments I, Root hair density of 7 -day-old seedlings. Root hair density was quantified in a 1 mm segment of the primary root 2 mm from the tip. Data are means of 3 independent biological replicates, >20 seedlings/plants in each. Bars with asterisks are significantly different from the mock (mean ± SD; ANOVA, P < 0.05, Tukey’s test).

### D27-LIKE1 is localised in plastids and displays a remarkable promoter activity in leaf midvein

To determine the subcellular localization of the D27-LIKE1 protein, we performed a transient expression experiment of D27-LIKE1 in L*er* protoplasts. *D27-LIKE1* cDNA was fused to the N terminus of GFP under the control of 35S promoter. As a positive control, a cytoplasm and nucleus-specific construct was used. D27 has been reported to be localised in plastids in rice (Hao *et al*., 2009) and in Arabidopsis (Waters et al., 2012). Consistent with these studies, GFP fluorescence fully overlapped with the red chloroplast autofluorescence (Fig. 4A). Thus, similar to its paralog, we can conclude that D27-LIKE1 resides in plastids.

**Figure 4.**
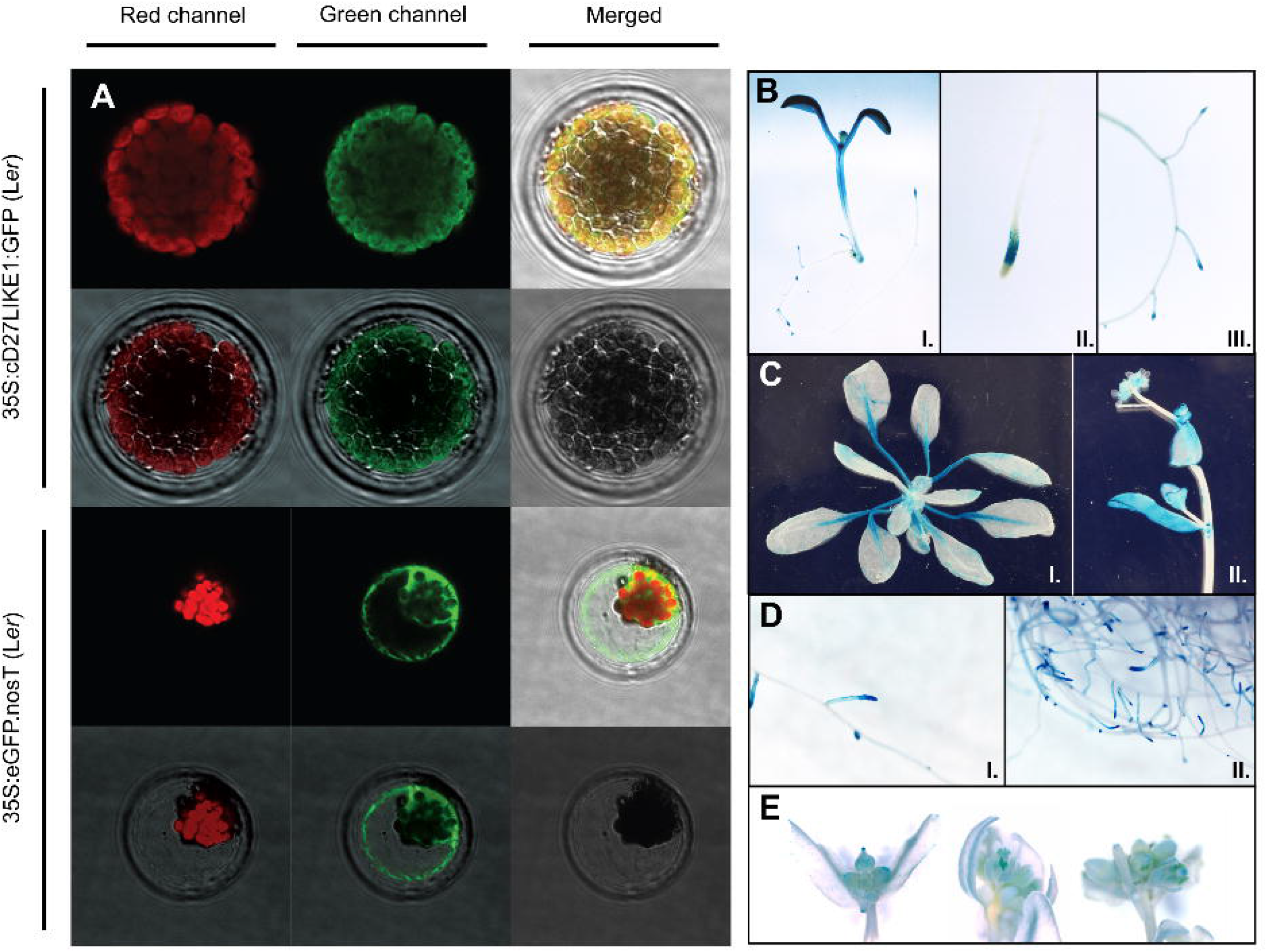
Subcellular localization of D27 -LIKE1 and tissue specific activity of the *D27-LIKE1* promoter. A, Chloroplast localization of 35S:cD27 -LIKE1:GFP in L*er* protoplast cells. Chlorophyll red autofluorescence and green fluorescence signals fully overlap on the merged image of 35S:cD27 -LIKE1:GFP (top panel) in contrast to 35S:eGFP:nosT (bottom panel). B-E, Spatio-temporal regulation of *D27-LIKE1* expression. *D27-LIKE1* promoter activity has been assessed using a 2.2 -kb promoter region of the gene fused to GUS coding sequence B, GUS activity in 7 -day-old seedlings in whole seedling (I.), in primary roots (II.) and in lateral roots (III.). C, GUS activity in 4-week-old rosette (I.), in stem and cauline leaves of 5-week-old plant (II.). D, GUS activity in 6-week-old roots. E, GUS histochemical activity in different stages of flower development.

To elucidate the spatio-temporal regulation of *D27-LIKE1* expression, *D27-LIKE1* promoter constructs fused with the GUS coding region has been generated. GUS expression was assayed in at least eight representative T_3_-T_5_ homozygous lines in three different developmental stages (Fig. 4B-E). 7-day-old seedlings displayed strong activity in the cotyledon, while expression was variable in the hypocotyl (Fig. 4B). Primary and lateral root tips showed a distinctive GUS signal in the elongation zone (Fig. 4B). GUS expression has also been assayed in 4-week-old rosettes (Fig. 4C). The *D27-LIKE1* promoter was found to be strongly active in the petiole and the midrib while the GUS signal faded away towards the leaf margin. In the reproductive plant stage, no signal was detected in the stem (Fig. 4C) and roots displayed the same expression pattern in the elongation zone as of young seedlings (Fig. 4D). Furthermore, we detected remarkable GUS staining also in the meristematic zone. In flowers, the stigma showed strong GUS activity, while sepals and stamens displayed low expression (Fig. 4E).

### *D27-LIKE1* mRNA is phloem mobile and transported to the root

In a comprehensive study, numerous mobile transcripts were identified in Arabidopsis phloem sap, where *D27-LIKE1* mRNA was found to be transported from shoot to root (Thieme *et al*., 2015). To prove that *D27-LIKE1* mRNA is indeed a phloem mobile signal, we collected phloem sap and analysed the extracted mRNA using the Droplet Digital PCR method (ddPCR), which is dedicated to detect very rare messages. As shown in Fig. 5A, the ddPCR analysis verified the presence of *D27-LIKE1* mRNA in phloem sap. To provide further evidence of *D27-LIKE1* mRNA mobility and to examine the proposed shoot-to-root transport, we generated grafts between L*er* scions and *d27-like1-1* rootstocks. Consistent with the previous findings, *D27-LIKE1* transcripts were present in the root tissues of *d27-like1-1* stocks two weeks after grafting (Fig. 5B-C), inferring that the *D27-LIKE1* transcript is phloem-mobile.

**Figure 5.**
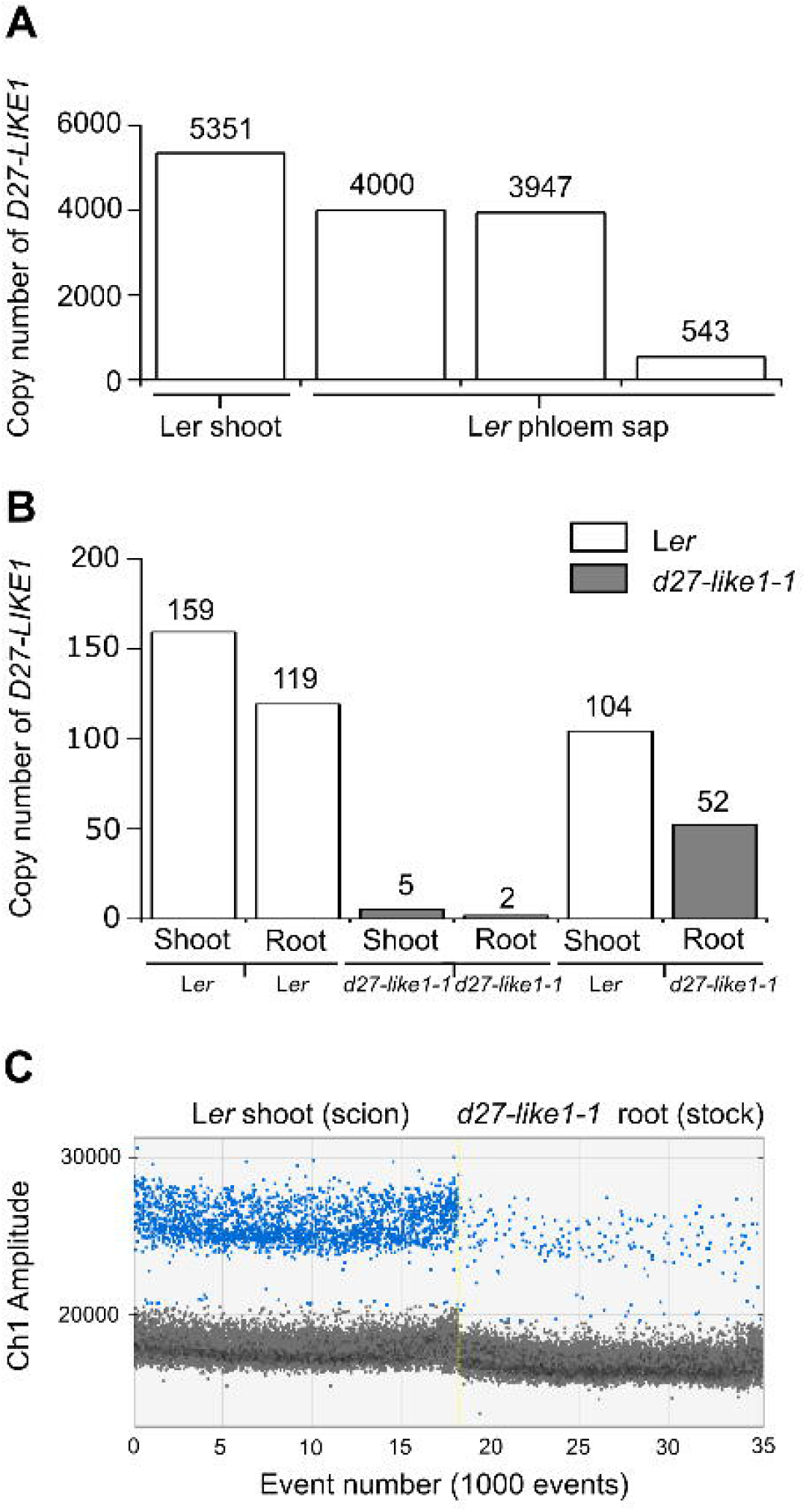
*D27-LIKE1* mRNA is phloem mobile and transported from shoot to root. A, Copy number validation of *D27-LIKE1* mRNA by droplet digital PCR (ddPCR) in shoot and in phloem sap collected in three experiments according to Tetyuk et al., 2013. Copy numbers were normalised and are expressed as per 1000 copy numbers of *UBQ10*. B, Copy number validation of *D27-LIKE1* mRNA by ddPCR in grafted seed lings (L*er*|L*er, d27-like1-1*|*d27-like1-1* and L*er*|*d27-like1-1)*. Copy numbers were normalised and are expressed as per 1000 copy numbers of *ACTIN2*. C, One dimensional plot of ddPCR assay showing the presence of *D27-LIKE1* mRNA in the roots of a L*er*|*d27-like1-1* grafted seedling. Positive droplets, containing the target DNA are shown with blue and negatives with gray colors. X-axis represents the total number of droplet events. Y-axis represents the detected fluorescence amplitude. Experiments were repeated three times (n=3).

### *d27-like1-1* does not influence the function of the photosynthetic apparatus

Carotenoids are essential in enhancing photosynthetic efficiency and preventing photodamage. To challenge photoprotective processes, chlorophyll fluorescence measurements were achieved in 14 d-old plants subjected to different light intensities for one week. The actual quantum yield [Y(II)], the heat dissipation process (NPQ) and the non-regulated heat dissipation process [Y(NO)] did not differ significantly in L*er* and *d27-like1-1* under all light regimes (Supplemental Figure S3). Thus, we can state that the disruption of *D27-LIKE1* has no noticeable effect on the function of the photosynthetic apparatus.

### The *d27-like1-1* mutant displays 9-*cis*-violaxanthin and 9-*cis*-β-carotene accumulation

As a close paralog of D27, D27-LIKE1 is predicted to be a putative β-carotene isomerase. To test whether the mutation of *D27-LIKE1* affected the carotenoid profile, wt, *d27-like1-1, D27-LIKE1* overexpressing and complemented lines were investigated in two parallel analytical systems. Numerous carotenoids were separated in *d27-like1-1* and L*er* plants (Fig. 6A-B, Supplementary Table S2). By overlapping the carotenoid profiles, it became clear that only one peak (6^th^) showed a massive, and another one (12^th^) displayed a slight increase in the mutants, which were identified as 9-*cis*-violaxanthin and 9-*cis*-β-carotene, respectively. *d27-like1-1* rosettes contained ∼3 times more 9-*cis*-violaxanthin and 30% more 9-*cis*-β-carotene than wt plants. To confirm that *d27-like1* mutation was solely responsible for this phenotype, we performed the carotenoid profiling with the complementing lines also. The wild-type level of 9-*cis*-violaxanthin was fully restored in all lines, while 9-*cis*-β-carotene level was partially complemented (Fig 6C-D), suggesting that the abruption of *D27-LIKE1* is responsible for these phenotypes. Intriguingly, *D27-LIKE1* overexpressing lines had similar carotenoid composition and profile as L*er* plants (Supplementary Table S2.).

**Figure 6.**
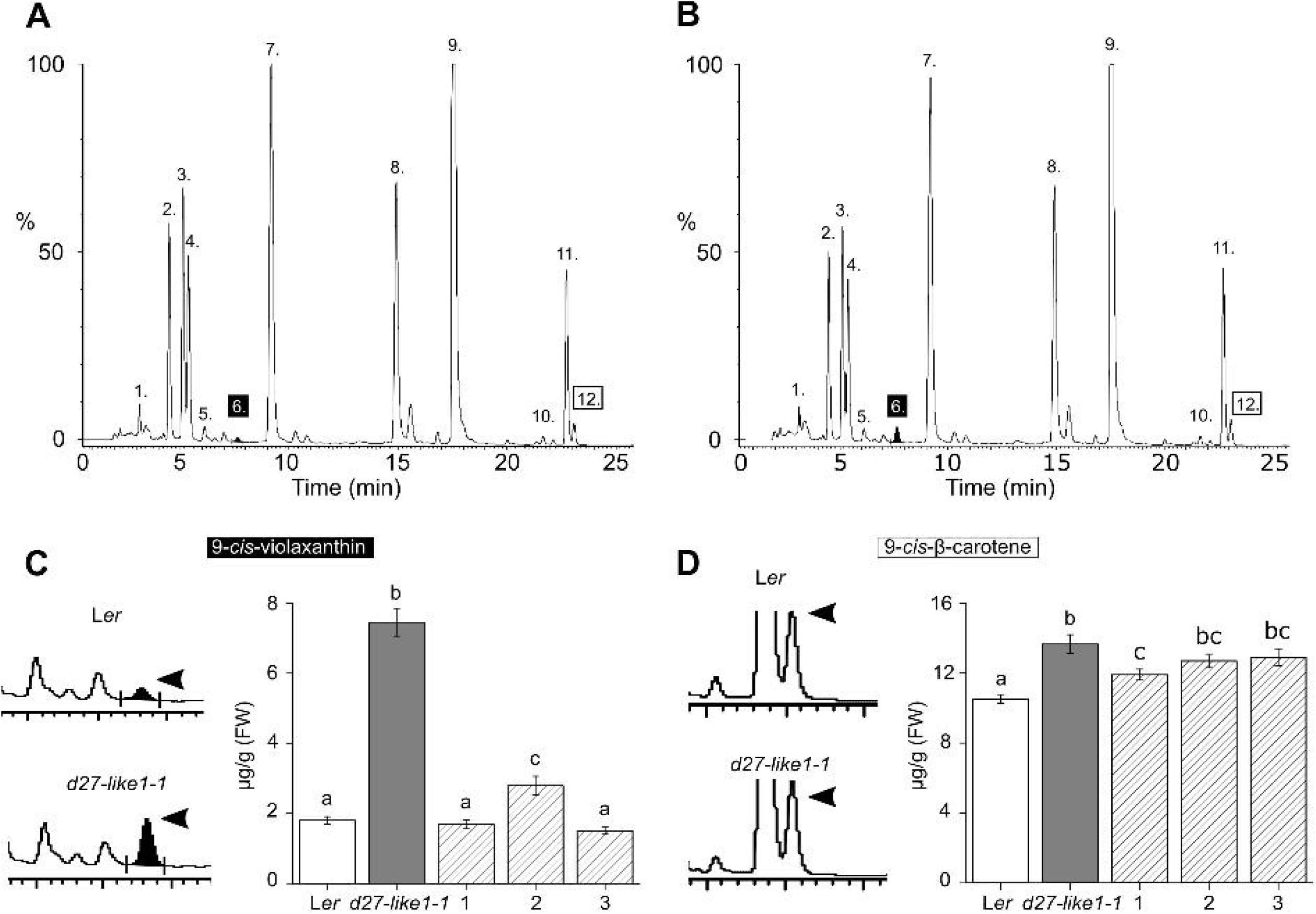
The *d27-like1-1* mutant accumulate 9-*cis*-violaxanthin. The carotenoid profile of 2 -week-old wild type (A) and *d27-like1-1* (B) rosettes has been inferred by HPLC. The levels are expressed as a percentage of the total carotenoid pool. Peak assignments: 1, chlorophyllide B; 2, chlorophyllide A; 3, *trans-*violaxanthin; 4, *trans-*neoxanthtin; 5, 13-*cis*-β-violaxanthin; 6, 9-*cis*-violaxanthin; 7, *trans*-lutein; 8, chlorophyll b; 9, chlorophyll a; 10, 13-*cis*-β-carotene; 11, all -*trans*-β-carotene; 12, 9 -*cis*-β-carotene. C (left) and D (left), Enlarged HPLC chromatograms of 9-*cis*-violaxanthin and 9-*cis*-β-carotene peaks. C (right) and D (right), 9 -*cis*-violaxanthin and 9-*cis*-β-carotene content of 2-week-old L*er, d27-like1-1* and three complemented mutant lines (denoted as 1, 2 and 3). Bars with the same letter are not significantly different from each other (mean ± SD; ANOVA, P < 0.05, Tukey’s test).

### Recombinant D27-LIKE1 catalyses *trans*/*cis, cis*/*cis* isomerisation of carotenoids *in vitro*

It has been previously shown that rice OsD27 and the AtD27 catalyse the reversible isomerization of 9-*cis*- and all-*trans*-β-carotene (Alder et al., 2012; Harrison et al., 2015; Abuauf et al., 2018). The high similarity of the consensus sequences prompted us to test whether D27-LIKE1 also acts as a carotenoid isomerase with similar substrate preferences. We expressed MBP-fused D27 and D27-LIKE1 enzymes without the predicted cTP in BL21(DE3) *E*.*coli* cells and performed *in vitro* assays with the purified proteins. Either with the cTP, or with a GST-tag, the recombinant proteins resided in the inclusion body. With the use of MBP-tag and a Fe-supplementation in growing media, the proteins became soluble and were produced with high yield. Purified recombinant proteins displayed a brownish colour, which indicates the presence of iron (Supplemental Figure S4).

Incubation of recombinant D27-LIKE1 protein with all-*trans*-β-carotene resulted in a detectable conversion into the major *cis* isomers of β-carotene (Fig. 7A; Supplementary Figure S5., Supplementary Table S3.). We observed a ∼4.5-fold increase in 9-*cis*-, ∼3.5-fold increase in 15-*cis*- and ∼2.5-fold increase in 13-*cis*-β-carotene compared to the control. As expected, D27 catalysed a more remarkable all-*trans*-β-carotene isomerisation into 9-cis-β-carotene, and intriguingly, towards 13-*cis*-β-carotene and 15-*cis*-β-carotene forms which was observed only *in vivo* previously (Abuauf *et al*., 2018).

**Figure 7.**
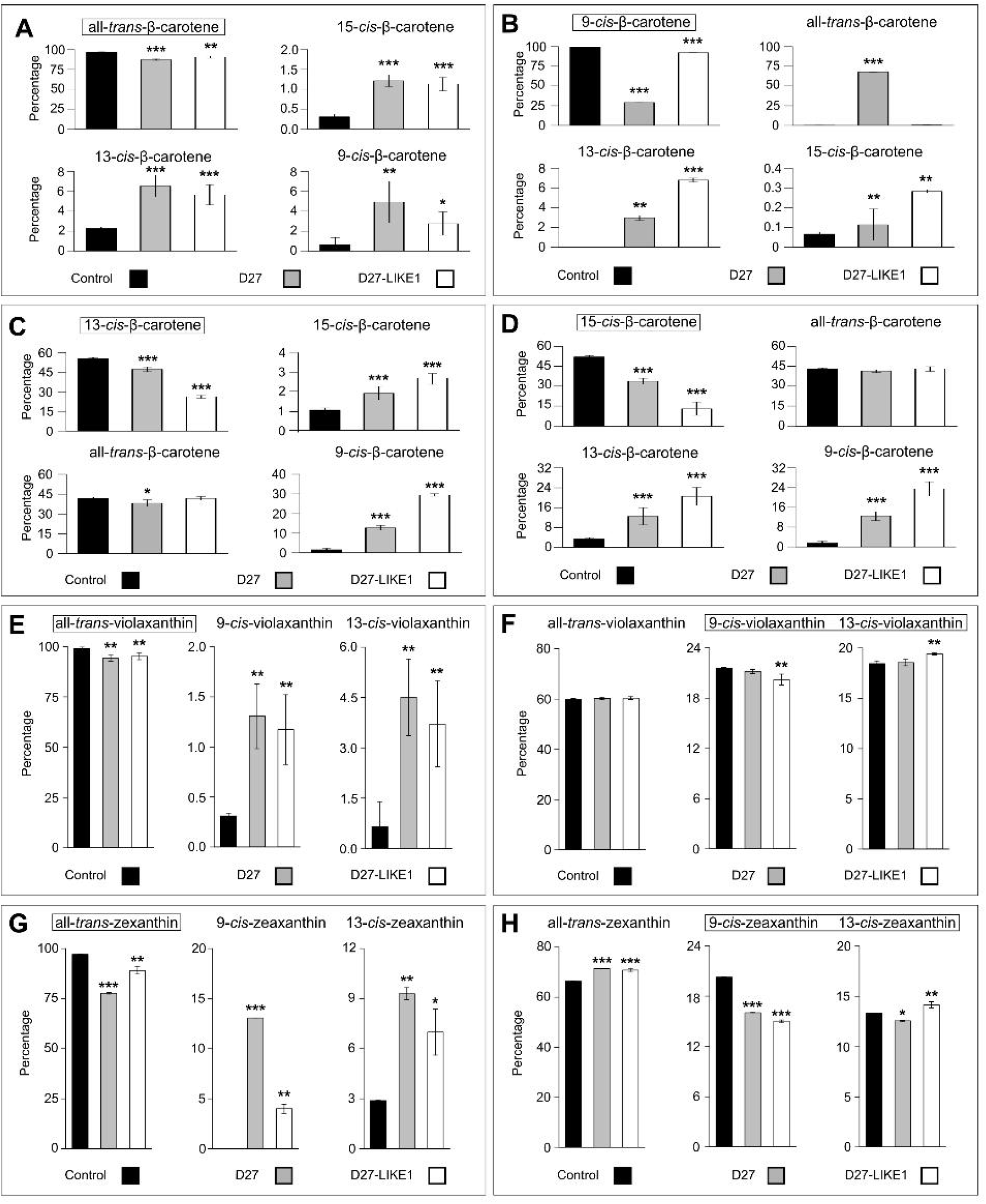
D27-LIKEl has a preferential trans -cis and cis -cis isomerase activity on carotene and xanthophyll substrates in vitro feeding experiments have been performed with recombinant D27-LIKE1, D27 and void plasmid construct (control) with various substrates. Percentage of different cis- and trans isomers (boxed names) when A, all-trans-ß-carotene; B, 9-cis-ß-carotene; C, 13-cis-ß-carotene: D, 15-cis-ß-carotene; E, all-trans-violaxanthin; F, mixture of cis -violaxanthin isomers (9 -cis, 13 -cis); G, all -trans-zeaxanthin; and H, mixture of cis -zeaxanthin isomers (9-cis, 13-cis) were incubated with 50 µg recombinant D27 or D27-LIKE1 protein. Data are means of four biological replicate experiments in each (n=4). Bars with asterisks are significantly different from the control (mean ± SD; ANOVA, *P < 0.05, **P < 0.01, ***P < 0.001, Tukey’s test).

Next, we tested whether the enzymes can catalyse the reverse *cis-trans* isomerisation and the interconversion between the *cis* forms. D27 very slightly isomerized 13-*cis*-β-carotene into all-*trans*-β-carotene and a massive isomerization from 9-*cis*-β-carotene into all-*trans*-β-carotene has been observed. Interestingly, D27-LIKE1 did not isomerise any of the *cis* forms into *trans* forms. Feeding with either 9-, 13- or 15-*cis*-β-carotene, the amount of either isomers increased, however, D27-LIKE1 had a more prominent activity regarding the interconversion between the *cis* forms (Fig. 7B-C; Supplementary Figure S5., Supplementary Table S3.). Both enzymes were capable to catalyse the interconversion between *cis/cis* forms, especially towards 9-*cis*-β-carotene, which was predominant in all cases tested. We can conclude that D27-LIKE1 is a preferential *trans/cis* and *cis/cis* isomerase while D27 is more capable to catalyse the reversible conversion between all-*trans*-β-carotene and 9-*cis*-β-carotene.

A ∼3-fold increase of 9-*cis*-violaxanthin content in *d27-like1* mutants exhorted us to test whether recombinant D27-LIKE1 can catalyze the isomerization of violaxanthin *in vitro*. Incubation with all-*trans*-violaxanthin resulted in 3.7-fold increase in 9-*cis*- and about 5.8-fold increase in 13-*cis*-violaxanthin amounts compared to the control (Fig. 7D; Supplementary Figure S5., Supplementary Table S3.). Intriguingly, recombinant D27 in our system also converted all-*trans*-violaxanthin into 9-*cis*- and 13-*cis*-violaxanthin. Interestingly, when a mixture of *trans*-*cis*-violaxanthin isomers containing 22% 9-*cis*- and 18% 13-*cis*-violaxanthin was added, no significant change has been observed in the amount of all-*trans*-violaxanthin suggesting that the enzymes did not convert *cis* to *trans* forms or a negative feedback mechanism prevented the *trans/cis* isomerization. (Fig. 7E; Supplementary Figure S5., Supplementary Table S3.). A slight decrease in 9-*cis*-violaxanthin and increase in 13-*cis*-violaxanthin amounts might be the consequence of 9-*cis*/13-*cis*-interconversion catalysed by D27-LIKE1 (Fig. 7F; Supplementary Figure S5., Supplementary Table S3.).

Next, we wanted to examine whether D27-LIKE1 isomerise zeaxanthin, a major component of the xanthophyll cycle and the first stable product of the all-*trans*-β-carotene hydroxylation. Incubation with all-*trans*-zeaxanthin led to a ∼4- and ∼2.5-fold increase in 9-*cis*- and 13-*cis*-zeaxanthin amounts (Fig. 7G; Supplementary Figure S5., Supplementary Table S3.), while incubation with D27 resulted in a more prominent formation of these *cis* zeaxanthin forms. To test the reverse isomerization and the possible feedback inhibition, a *cis*-zeaxanthin mixture with 20% 9-*cis*- and 13% 13-*cis*-zeaxanthin composition was added as a substrate (Fig. 7H; Supplementary Figure S5., Supplementary Table S3.). Compared to the control we detected a slight, albeit significant increase in the amount of all-*trans*-zeaxanthin with both enzymes suggesting that a *cis/trans* isomerisation is also possible with zeaxanthin substrates. The assay revealed a significant decrease in 9-*cis* and a slight increase in 13-*cis* isomeric forms similarly to the *cis*-violaxanthin mixture test (Fig. 7E, Supplementary Figure 5., Supplementary Table S3.).

### *d27-like1-1* mutant displays a moderate increase in ABA and PA content

Higher levels of the ABA biosynthesis precursor 9-*cis*-violaxanthin are associated with higher levels of ABA (Neuman *et al*., 2014). The carotenoid profile of *d27-like1-1* plants shows remarkable 9-*cis*-violaxanthin accumulation, which might eventually results in a higher ABA content. To test this hypothesis, we measured ABA and phaseic acid (PA; Kushiro et al., 2004) levels in rosettes of 14d old *d27-like1-1* and L*er* plants. Both compounds displayed a slight, ∼1.7 - 1.9-fold increase in *d27-like1-1* plants (Fig. 8A). Next, to provoke stress-induced ABA accumulation, we assessed the ABA and PA content of 14d-old detached rosettes at the stage when they reached 20% water loss. As expected, dehydration resulted in ABA accumulation in both genotypes, however, ABA levels in stressed *d27-like1-1* rosettes did not exceed wt levels (Supplementary Figure S6.).

**Figure 8.**
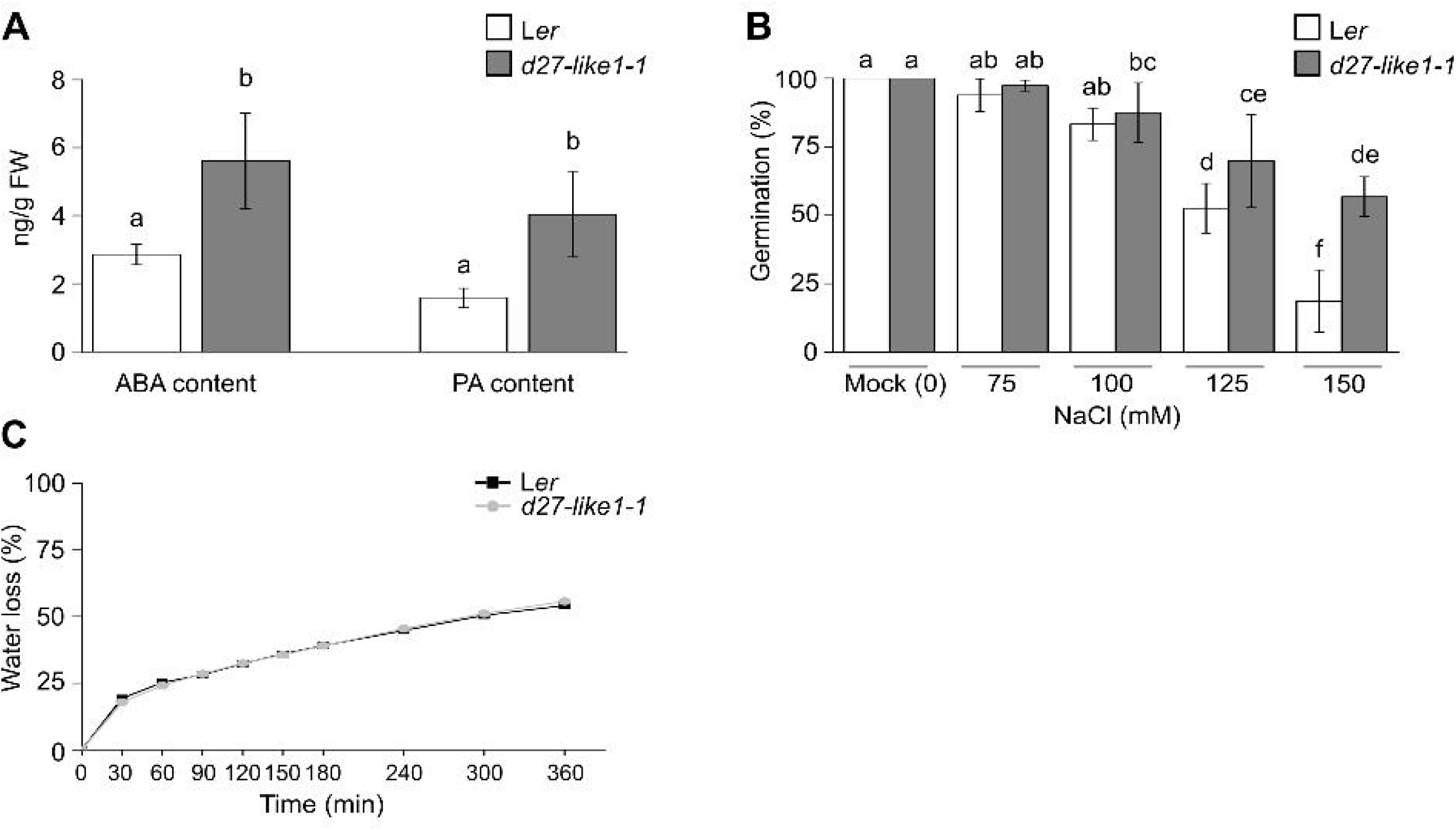
*d27-like1-1* mutant exhibits a moderate increase in ABA and PA content ABA and PA levels as well as the salt stress and dehydration tolerance of *d27-like1-1* has been investigated. A, ABA and PA content of 2 -week-old L*er* and *d27-like1-1* detached rosettes. Data are means of 5 independent biological replicates, 5 rosettes per genotype in each. B, Germination efficiency of *d27-like1-1* on NaC1 supplemented plates Germination percentages were assessed in 7 -day-old seedlings. Data are means of 3 independent biological replicate experiments, >60 seedlings per genotype in each. Bars with the same letter are not significantly different from each other (mean ± SD; ANOVA, P < 0.05, Tukey’s test). C, Water loss of detached L*er* and *d27-like1-1* rosettes over a 6hrs long period. Data are means of 5 independent biological replicate experiments, 5 rosettes per genotype in each (mean ± SD; ANOVA, P < 0.05, Tukcy’s test)

Genotypes with constitutively elevated ABA level display slight-to-moderate stress resistance phenotype. Detached rosettes of *AtNCED6* overexpressing Arabidopsis lines are slightly resistant to water loss (Finn *et al*., 2015) while *AtNCED3* overexpressing lines exhibit lower transpiration rates (Iuchi *et al*., 2001). Similarly, *ABA2* overexpressing Arabidopsis lines are more tolerant to salinity when grown on agar plates or soil (Lin *et al*., 2007). To prove that the slightly elevated ABA level of *d27-like1-1* plants is accompanied by a higher tolerance of water loss and salinity stress, we performed experiments with detached rosettes as well as with seeds germinating under different salinity regimes. We found that *d27-like1-1* plants did not show a significantly different, decreased water loss under our experimental conditions (Fig. 8C). However, *d27-like1-1* seeds displayed a significantly better germination under higher salt concentrations (Fig. 8B). The overall germination capability of both genotypes has been affected as a consequence of high salinity, as previously described ((Lin et al., 2007). Lower concentrations of NaCl (75, 100 mM) had no notable effect on the germination percentage, however, significant differences were observed on higher salt concentrations. In the presence of 125 mM NaCl, *d27-like1-1* exhibited better germination percentages than L*er*. This difference increased further when a concentration of 150 mM was applied, where the germination percentage of *d27-like1-1* seeds was approximately three times higher than wt seeds. Taken all round, these results indicate that the slightly elevated constitutive ABA level of *d27-like1-1* plants is not necessarily manifested in higher stress tolerance, albeit the salinity

## Discussion

### D27-LIKE1 is a preferential *trans/cis* and *cis/cis* isomerase of β-carotene substrates *in vitro*

D27 has been in focus for years and its enzymatic characteristics have been extensively studied (Alder et al., 2012; Harrison et al., 2015; Bruno and Al-Babili, 2016; Abuauf et al., 2018). *In vivo*, D27 converts all-*trans*-β-carotene to *cis*-β-carotene isomers including 9-, 13- and 15-*cis*-β-carotene with a pronounced 9-*cis*-β-carotene preference (Abuauf *et al*., 2018). However, recombinant D27 did not convert all-*trans*-β-carotene into 13- or 15-*cis*-β-carotene (Abuauf *et al*., 2018). In contrast, we detected a moderate 13- and 15-*cis*-β-carotene accumulation under our experimental conditions when recombinant D27 was fed with all-*trans*-β-carotene. We observed a slight, *cis/trans* conversion to all-*trans*-β-carotene from 13-*cis*-β-carotene and found that D27 is capable to catalyse the interconversion between the *cis* forms. Abuauf et al. (2018) also found that feeding recombinant D27 with 9-*cis*-β-carotene yield not only the accumulation of the *trans* configuration but 13- and 15-*cis*-β-carotene isomers as well, however, incubation with 13- and 15-*cis* isomers did not result in interconversion. A similar specificity has been demonstrated in a rice D27 system except that 13-*cis*-β-carotene has not been detected (Bruno and Al-Babili, 2016).

Recombinant D27-LIKE1 performed very similarly to D27 in the *in vitro* feeding experiments. It converted all-*trans*-β-carotene into 9-, 13- and 15-*cis*-β-carotene although D27 produced remarkably higher amount of 9-*cis*-β-carotene. On the contrary, the enzyme interconverted the *cis* configurations of β-carotene to other stereoisomers in higher quantities than D27 proposing that D27-LIKE1 exhibits a more efficient *cis*-*cis* activity. Intriguingly, D27-LIKE1 did not convert any of the *cis* configurations to the *trans* isomer, suggesting that D27-LIKE1 is a preferential *trans*/*cis, cis*/*cis* isomerase. In plants, 9-, 13- and 15-*cis* isomers of β-carotene represent a minor, 5-30% portion of the total β-carotene pool and contribute to the colour of fruits and vegetables (Khoo *et al*., 2011). 13- and 15-*cis* forms have not been assigned as precursors of any known apocarotenoids yet. All-*trans*-β-carotene is isomerized non-enzymatically to *cis* forms when exposed to heat or by photoexcitation (Guo, Tu and Hu, 2008). Of the *cis* forms, 13-*cis*-β-carotene formation is kinetically favoured (Guo, Tu and Hu, 2008), which is in good agreement with its relative abundance among other *cis* isoforms. In plastids and solvents, 9-*cis* and 13-*cis* isomers were found to be the predominant isomers thus all-*trans*-β-carotene is relegated as the reservoir of the *cis* forms (Aman, Schieber and Carle, 2005). We propose that D27-LIKE1 is responsible for maintaining an optimal physiological concentration of 9-*cis*-β-carotene when conditions permit the non-enzymatically and uncontrolled isomerization of *trans* and *cis* forms (Fig. 9).

**Figure 9.**
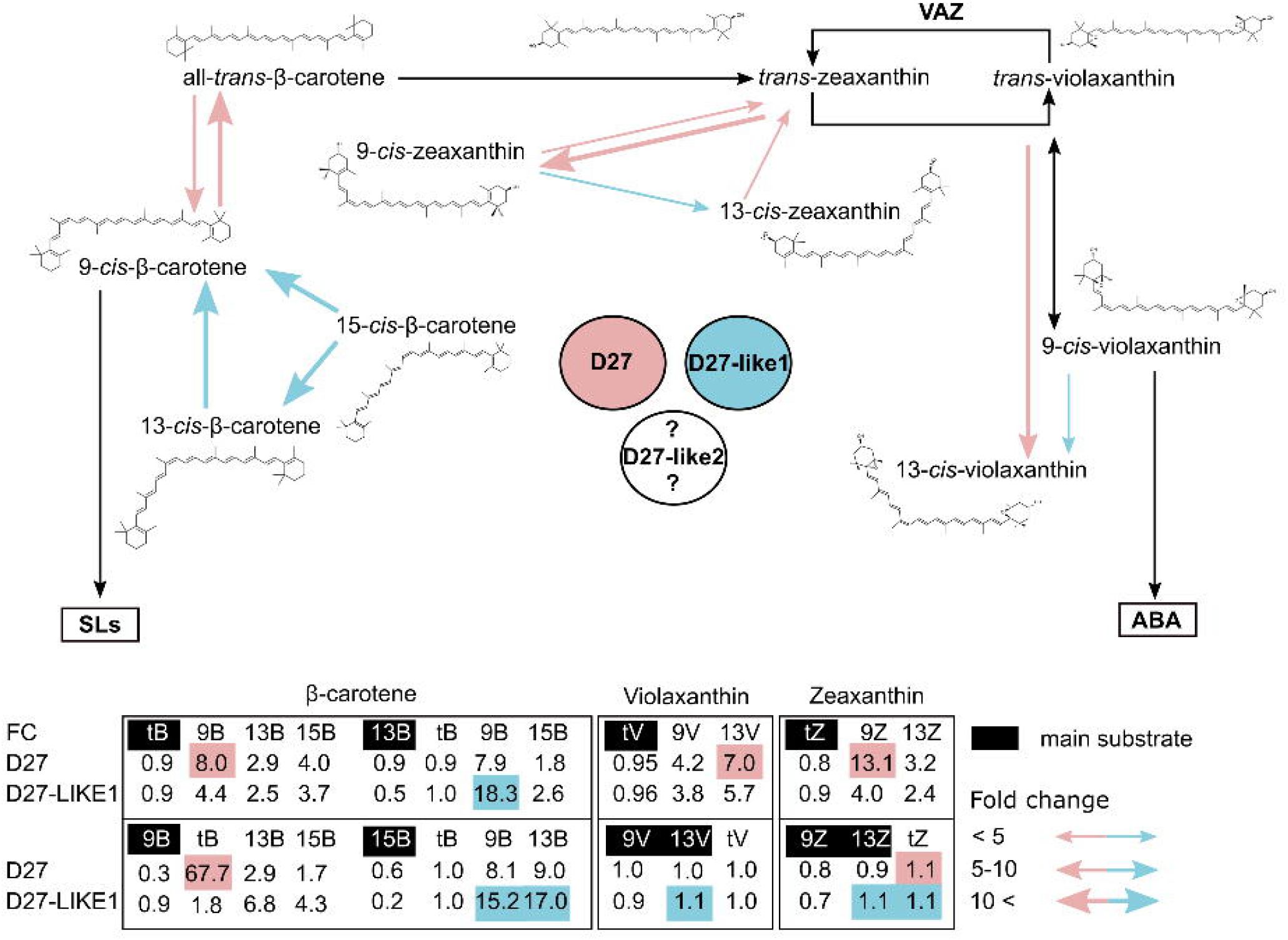
Working scheme of D27 and D27 -LIKE1-mediated isomerization of carotene and xanthophyll substrates in Arabidopsis. D27-LIKE1 is a *trans-cis*, and - more preferentially - a *cis-cis* isomerase of 9-, 13- and 15-*cis*-β-carotene forms and responsible for maintaining an optimal physiological concentration of 9 *-cis-*β*-*carotene when the non -enzymatical and uncontrolled isomerization of *trans* and *cis* forms is favoured. On the contrary, D27 is more capable to catalyse the reversible conversion between all *-trans-*β*-*carotene and 9 *-cis-*β-carotene and provides the precursor 9-*cis*-β-carotene for the SL biosynthesis *in planta*. Both D27 and D27-LIKE1 are capable to convert *trans*-violaxanthin and *trans*-zeaxanthin to 9- and 13*-cis* forms *in vitro*. Only D27-LIKE1 can catalyse the interconversion between the 9 - and 13 -*cis*-violaxanthin. Furthermore, *d27-like1-1* mutant accumulates 9-*cis*-violaxanthin, which is one of the known ABA precursors. Taken together, we propose that D27-LIKE1 is primarily a violaxanthin isomerase *in planta* which keeps the balance between *trans* and *cis* isomer configurations, and in the second place, can act as an auxiliary 9 -*cis*-β-carotene isomerase which produces precursor for the SL biosynthesis. The precise modulation of the enzymatic activity of the two enzymes by expression regulation, phloem trans port or imputs from the carotenoid flux eventually lead to a finely adjusted *cis/trans* ratio and eventually, an optimised apocarotenoid precursor availability. The table shows the fold changes in feeding experiments in which D27 (pink) and D27-LIKE1 (blue) recombinant protein were fed with various substrates *in vitro* (tB: *trans*-β-carotene, 9B; 9 -*cis*-β-carotene, 13B: -*cis*-β-carotene, 15B: 15 -*cis*-β-carotene, tV: *trans*-violaxanthin, 9V: 9 -*cis*-violaxanthin, 13V: 13 -*cis*-violaxanthin, tZ: *trans*-zeaxanthin, 9Z: 9-*cis*-zeaxanthin, 13Z: 13 -*cis*-zeaxanthin). Carotenoids highlighted in black are the main substrates in assays. The values highlighted in pink and blue are the most significant fold changes in feeding assays. The thickness of arrows indicates the rate or fold changes (<5, 5-10, 10<). VAZ: Xanthophyll cycle.

The capability of D27-LIKE1 to convert a minute amount of all-*trans*-β-carotene to 9-*cis*-β-carotene and a massive interconversion of 13- and 15-*cis* forms into 9-*cis* isomer raises the question whether it is an auxiliary, *bona fide* SL biosynthesis enzyme acting redundantly with D27. Clearly, *d27-like1-1* did not exhibit any SL/karrikin deficiency-related phenotype and indistinguishable from wt. Conversely, Arabidopsis *d27* displays a mild branching phenotype compared to the *d14* signalling mutants (Waters et al., 2012) suggesting that an alternative enzyme performs the isomerisation of all-*trans*-β-carotene. We propose that D27-LIKE1 might contribute to the SL biosynthesis *in planta*; however, its involvement is overwhelmed by D27, which is a more potent enzyme of all-*trans*-β-carotene conversion into 9-*cis*-β-carotene. D27-LIKE1 has no *cis/trans* isomerase activity and preferentially catalyses the interconversion of *cis* stereoisomers, which are less abundant in plant tissues, therefore it has a substantially lower substrate input flux to produce the direct precursor of SL biosynthesis (Fig. 10). To fully resolve the potential involvement of D27-LIKE1 in SL biosynthesis, higher order mutants with *d27, d27-like1, and d27-like2* should be generated in the future, which are expected to display a stronger branching phenotype than *d27*.

### D27-LIKE1 has a primary role in setting the ABA precursor 9-*cis*-violaxanthin levels *in planta*

In a seminal publication the substrate specificity of D27 has been assessed and the authors found that D27 did not convert neither zeaxanthin, nor violaxanthin, however, accepts α-carotene and all-*trans*-β,β-cryptoxanthin as substrate (Bruno and Al-Babili, 2016). Intriguingly, both D27-LIKE1 and D27 took the xanthophylls violaxanthin and zeaxanthin as substrate in our experimental system and produced a minute, albeit significantly higher amount of *cis* forms *in vitro*. We speculate that the higher sensitivity and the better selectivity of our UPLC detection system or minor differences between the experimental conditions might have led to this discrepancy. D27 contains iron (Lin *et al*., 2009), which is quintessential for the enzymatic activity (Harrison *et al*., 2015). Recombinant D27-LIKE1 protein stocks had greyish brown colour, which is a good indicative of the presence of iron (Lin *et al*., 2009). For the optimal formation of iron-sulphur clusters, the background Fe and S content of the growing media is sometimes not sufficient (Jaganaman et al., 2007; Nakamura et al., 1999), therefore, supplementation of the induction media with a mixture of iron forms in various oxidation states and cysteine is recommended (Jaganaman *et al*., 2007). We believe that this additional step contributed to a better catalytic activity of our recombinant protein stocks.

Optimal ABA biosynthesis requires ABA4 and NXD1 and occurs preferentially via 9-*cis*-neoxanthin, rather than 9-*cis*-violaxanthin (Perreau *et al*., 2020), which consists of only ∼6% of the violaxanthin pool in Arabidopsis. Both *aba4* (North *et al*., 2007) and *nxd1* (Perreau *et al*., 2020) Arabidopsis plants display high *trans*-violaxanthin and 9-*cis*-violaxanthin levels, with concomitant higher ABA level observed only in the *nxd1* mutant (Perreau *et al*., 2020). ABA4, in the absence of functional NXD1, is capable of promoting the isomerization of *trans*-violaxanthin. Interestingly, neither ABA4 nor NXD1 have direct *trans*/*cis* isomerase activity, hence require an unknown cofactor or additional isomerases (Perreau *et al*., 2020). We presented evidence that *d27-like1-1* accumulates high level of 9-*cis*-violaxanthin and both D27 and D27-LIKE1 can convert *trans*-violaxanthin to a minute amount of 9-*cis*-violaxanthin *in vitro*, therefore, they might be good candidates for the long-sought subtle isomerase, which produce one of the known ABA precursors. Rice *d27* mutants display reduced drought tolerance (Haider *et al*., 2018), cold survival rate (Liu *et al*., 2020) and lower ABA levels when compared to wt plants (Haider *et al*., 2018; Liu *et al*., 2020). Interestingly enough, these mutants have slightly reduced violaxanthin/neoxanthin content, while rice *D27* overexpressing lines exhibit higher ABA, violaxanthin/neoxanthin and lutein levels (Haider *et al*., 2018). These findings indicate that D27 somehow interferes with ABA biosynthesis, although it can’t be ruled out that D27 affects ABA metabolism through SL signalling, as observed in Arabidopsis *d14* plants (Li *et al*., 2020). Similar to rice *d27* plants, shoots of Arabidopsis *d27* mutant contain ∼20% less ABA than wt (Abuauf *et al*., 2018). Nevertheless, *d17* and *d10* SL mutants of rice (corresponding to Arabidopsis *MAX3* and *MAX4*, respectively) have a contrasting, higher ABA levels than *d27* (Haider *et al*., 2018), suggesting that D27 directly links ABA and SL biosynthesis *per se*. All in all, the enzymatic involvement of D27 in the ABA biosynthesis precursor is less understood and definitely requires additional research.

Contrary to Arabidopsis and rice *d27* mutants, *d27-like1-1* plants display slightly higher ABA content than wt plants which might be the direct consequence of 9-*cis*-violaxanthin accumulation, as observed in the *nxd1* mutant (Perreau *et al*., 2020). It has to be noted that *nxd1* accumulated not only the *cis*, but the *trans* isomer of violaxanthin and exhibited a remarkable, 2,5-times higher ABA content in detached rosettes than wt (Perreau *et al*., 2020). Intriguingly, rice *d27-like1* and *2* mutant plants exhibit lower ABA levels, similar to *d27* (Liu *et al*., 2020). It is tempting to speculate that in Arabidopsis, D27-LIKE1 acts primarily as a violaxanthin isomerase *in planta*, which keeps the balance between *trans* and *cis* isomer configurations, hence the high level of 9-*cis*-violaxanthin in the mutant plants. The *in vitro* feeding experiments, where D27-LIKE1 catalysed the *trans/cis* and *cis*/*cis* conversion, strongly support this notion.

### Functional divergence in the DWARF27 family

A higher resolution phylogenetic analysis of *DWARF27* ancestry revealed that although the previously suggested distinction of D27-like proteins is largely consistent with our results, we identified a fourth member of the family, *D27-LIKE3*, which is a remnant of an ancient duplication event and lost when flowering plants emerged. Our tree topology also suggests that while *D27-LIKE2* sequences can be found in all land plants, *D27-LIKE1* and *D27* are sister branches and are absent from certain taxonomic categories. It has been suggested that although SL synthesis is an ancient feature in land plants, SL signalling via canonical D14-type SL receptors is a relatively recent innovation in seed plants (Walker *et al*., 2019). Although a D14-like protein is present in Gymnosperms, the question arises whether a non-canonical SL signalling exist in the absence of the SMXL7 clade. If a canonical SL biosynthesis is operational in Gymnosperms, then the intriguing question is whether D27-LIKE1 might function as a major enzyme in the absence of D27. We demonstrated that D27-LIKE1 could catalyse the *trans*/*cis* conversion of all-*trans*-β-carotene *in vitro*. Based on this evidence we can propose that - if no alternative pathway exists - Gymnosperm D27-LIKE1 might be responsible for 9-*cis*-β-carotene production, although this prediction should be confirmed with experimental data. Nevertheless, the fact that either *D27* or *D27-LIKE1* are easily lost in several taxa implies that the enzymes might function redundantly and can replace each other to some extent. From a phylogenetic point of view, it is intriguing to consider that both enzymes have a parallel evolutionary history, which suggests that their enzymatic features might have been separated early and been chiselled to different functions. Although their broad substrate specificity is superimposable quite well, their isomer conversion preferences are different *in vitro*. Recombinant D27-LIKE1 accepts quite few substrates which indicates that the enzyme has a broad specificity with a characteristic preference towards *trans*/*cis* and *cis*/*cis* conversions of C_40_ carotenoid and xanthophyll substrates. The functional divergence of the enzymes is more apparent when their physiological role is investigated. D27 is evidently connected to SL synthesis, while D27-LIKE1 has no clear function in the SL/KL context with a weak, auxiliary 9-*cis*-β-carotene production from *trans*-β-carotene and a remarkable interconversion of 13- and 15-*cis* to 9-*cis* forms. On the contrary, *d27* rice and Arabidopsis plants have lower ABA levels than wt, while D27-LIKE1 adjusts 9-*cis*-violaxanthin content *in planta*, which is eventually manifested in the slightly elevated ABA level in the mutant. Besides, the overlapping substrate specificity and the spectrum of mild - to no apparent phenotypes indicate that the enzymes act partially redundantly *in vivo* (Fig. 9). A functional redundancy might explain the mild branching phenotype of *d27* Arabidopsis plants (Waters et al., 2012) and why *D27-LIKE1* overexpressing lines has no detectable alterations in their carotenoid profile.

D27-LIKE1 resides in the plastids similarly to Arabidopsis and rice D27 (Lin et al., 2009; Waters et al., 2012) along with other SL biosynthesis enzymes (Booker et al., 2004) favouring carotenoid metabolic channelling. In addition, the early steps of ABA synthesis and the synthesis of the VAZ cycle xanthophylls take place in the plastids. We provided evidence that *d27-like1-1* plants have a normal, functional photosynthetic apparatus indicating that the VAZ cycle is not perturbed. The tissue-specific activity of the *D27-LIKE1* promoter partially overlaps with the expression pattern of D27. In Arabidopsis, *D27* is expressed mostly in lateral root tips (Abuauf *et al*., 2018), which is consistent with the role of SLs in inhibiting lateral root growth (Rasmussen et al., 2012). On the contrary, D27-LIKE1 promoter is active in the lateral and primary root tip, raising the question whether it can mediate all-*trans*-β-carotene conversion in tissues where *D27* is less abundant or not expressed. Another dimension of *D27-LIKE1* expression is that the mRNA is phloem mobile and therefore could act in tissues where the *D27-LIKE1* promoter is not active. We could not detect GUS reporter activity in the vascular tissues of the root elongation and maturation zones; however, given that *D27-LIKE1* is a mobile signal, the mRNA can be translated here. It is noteworthy that *D27* mRNA has not been assigned as a phloem-mobile signal, while *D27-LIKE2* mRNA, along with *D27-LIKE1* has been transported in the phloem (Thieme *et al*., 2015).

Finally, we propose that a finely adjusted expression pattern, phloem transport or a precise setting of the molecular environment (for instance, input from the carotenoid flux, redox status, *cis*/*trans* ratio of respective carotenoids, spontaneous isomerisation etc.), might largely modulate the outcome of the enzymatic functions of the two enzymes. These events along with the enzymatic activity of the two proteins might eventually lead to a precisely balanced isomer constitution and hence, an optimised apocarotenoid precursor availability (Fig. 9).

## Materials and methods

### Phylogenetic Analysis

Protein sequences with homology to Arabidopsis D27 family members were identified by BLASTP searches of GenBank protein (https://www.ncbi.nlm.nih.gov/) and ONEKP (https://db.cngb.org/onekp/) transcriptome databases. Over 700 sequences were identified with highly significant E-value hits (<10^−20^). Alignments were obtained using MUSCLE with default settings (Edgar, 2004). The alignment was conservatively edited using PFAAT (http://pfaat.sourceforge.net) to remove regions of poor homology, then the aligment has been manually edited in Mega6 (Tamura *et al*., 2013). In total, 578 aligned sequences were used for the analyses (Supplementary Table S1). The protein sequences of the D27 family in Arabidopsis were subjected to domain prediction by InterPro search (Blum *et al*., 2021). The HMMER algorithm identified the 85 aa long DUF4033 domain which was extracted from the downloaded sequences. The phylogenetic tree has been constructed using the DUF4033 sequences and created in Geneious Prime 2021.2.2 (https://www.geneious.com/) with the NJ method with default settings. *Chlamydomonas reinhardtii* served as outgroup. The motif logos were created using the WebLogo tool (Crooks *et al*., 2004).

### Plant material and growth conditions

Seeds of *d27-like1* mutants (GT13552, GT22003) were obtained from the Cold Spring Harbor Laboratory (Cold Spring Harbor, NY, USA). Mutants were backcrossed three times to L*er*, and homozygous plants were selected by PCR with primers attb.D27-l1prom.F - D27-l1.R/Ds5-1 and D27-l1.cds.F/D27-l1.cds.R (all primers are listed in Supplementary Table S4.). PCR amplifications were accomplished with Phusion DNA Polymerase (Thermo Fisher, Waltham, MA, USA). Mutant line was named *d27-like1-1*. Landsberg *erecta* (L*er*) ecotype was used as wild-type (wt). For phenotyping, chlorophyll fluorescence analysis, carotenoid/hormone profiling and water loss experiments, plants were sown in Compo-Sana soil mixture (Compo, Münster, Germany). Plants were grown in Conviron or SANYO environmental chambers with 16 h/8 h or 8 h/16 h photoperiod (100 µmol m^−2^ s^−1^, 21°C/18°C, 75% relative humidity) unless otherwise stated. The chambers were equipped with Valoya L14 LED light sources (Valoya, Helsinki, Finland). For GUS assay, hypocotyl elongation assay and root hair density assay surface sterilised seeds were stratified at 4°C in the dark for 3 days and then sown on ½ MS medium containing 10 gL^-1^ sucrose and supplemented with 1 µM *rac*-GR24 (OlChemIm, Olomouc, Czech Republic) or 1 µM KAR2 (OlChemIm, Olomouc, Czech Republic), if necessary. For salt stress treatments sterilised and stratified seeds were sown on ½ MS medium containing 10 gL^-1^ sucrose and 75, 100, 125 or 150 mM NaCl. All Petri dishes were placed in chambers with 16 h/8 h photoperiod under the same environmental conditions as mentioned above.

### Overexpression and complementation of *d27-like1-1*

To generate overexpressing lines (*p35S:cD27-LIKE1*), *D27-LIKE1* cDNA was amplified using primers attb.D27-l1.cds.F/attb.D27-l1.cds.stop.R, then cloned into pDONR221 donor vector with BP clonase (Thermo Fisher, Waltham, MA, USA) and recombined into pGWB502 (Nakagawa *et al*., 2007) with LR clonase (Thermo Fisher, Waltham, MA, USA). The expression of *D27-LIKE1* has been assessed in homozygous overexpressing lines (Supplementary Figure S7.)

To generate complementation lines (*pD27-LIKE1:cD27-LIKE1* in *d27-like1-1*), 2.2 kb promoter fragments (with 5’UTR) were amplified with primers attb.D27-like1.prom.F/attb.D27-like1.prom.R and cloned into the BamHI/EcoRI site of pENTR4 vector (Thermo Fisher, Waltham, MA, USA). *D27-LIKE1* cDNA has been cloned into the EcoRI/NotI site of pENTR4 harbouring the *D27-LIKE1* promoter fragment. The assembled construct has been recombined into pGWB501 (Nakagawa *et al*., 2007). Vectors were introduced into *Agrobacterium tumefaciens* GV3101 and used for floral dip transformation (Végh et al., 2017). T_3_-T_5_ homozygous transgenic plants were used in all experiments.

### RNA Extraction and qPCR Analysis

Total RNA was isolated and DNAse I digested using RNEasy Plant Mini Kit (Qiagen, Hilden, Germany) according to the manufacturer’s instructions. cDNA was reverse transcribed with M-MuLV reverse transcriptase (Promega, USA). The qRT-PCR analysis was performed on an Applied Biosystems Fast7500 device using Fast SYBRGreen Master Mix (Applied Biosystems, Waltham, MA, USA). For each biological replicate, three technical replicates were examined. Arabidopsis *ACTIN2* was used as an endogenous control and the relative changes in gene expression were analysed according to the 2^−ΔΔCt^ method.

### GUS Histochemical Assay

To generate *pD27-LIKE1:GUS*, the 2.2kbp long promoter of *D27-LIKE1* was amplified with the primers attb.D27-like1.prom.F/attb.D27-like1.prom.R, then cloned into pDONR221 and recombined into pGWB533 (Nakagawa *et al*., 2007). Plants were grown either for 7 or for 28 days on ½ MS or for 21 or 28 days on soil. GUS histochemical staining was performed as described by Végh et al., 2017. Samples were incubated in the staining solution for 2-3 h at 37*°C* in the dark. Photographs were taken with a Leica DMS3000 microscope.

### Subcellular Localization

To generate *35S:cD27-LIKE1:GFP*, cDNA was amplified with the primers attb.D27-like1cDNA.F/attb.D27-like1cDNAnostop.R. Positive clones of pDONR221 were recombined into pGWB405 (Nakagawa *et al*., 2007). The resulting plasmid was introduced with the control plasmid 35S:eGFP:nosT into L*er* leaf protoplast as described (Wu *et al*., 2009). After overnight incubation in the dark, GFP signal and chlorophyll autofluorescence were detected with a Leica TCS-SP8 confocal microscope with the same exposure parameters at the excitation wavelength of 488 *nm*.

### Phloem Exudate and ddPCR Analysis

Phloem exudates were collected from 5 week-old L*er* rosettes as previously described (Tetyuk, Benning and Hoffmann-Benning, 2013). 100 U RNase inhibitor (Thermo Fisher, Waltham, MA, USA) was added to 1 ml of collected samples. Phloem cDNA analyses were performed on a QX200 Droplet Digital PCR (Bio-Rad, Hercules, CA, USA) according to the manufacturer’s instructions using endogenous control primers ACT2.F/ACT2.R and UBQ10.F/UBQ10.R, negative control primers rbcL.F/rbcL.R and *D27-LIKE1* primers D27-like1.RT.F/D27-like1.RT.R. The fluorescence intensity of the individual droplets was quantified using the QX200 Droplet Reader (Bio-Rad, Hercules, CA, USA) and analysed with QuantaSoft software (Bio-Rad, Hercules, CA, USA).

For grafting L*er* and *d27-like1-1*, seedlings were germinated on ½ MS for 7 days. Root-shoot grafts were fixed with silicone tubes (0,6 mm internal diameter) as described (Bainbridge *et al*., 2013). Grafted seedlings were grown on ½ MS for further 2 weeks before shoot and root samples were collected. Adventitious roots were regularly removed, and both scion and stock samples were genotyped. RNA samples were isolated from individual grafted plants and reverse transcribed for ddPCR.

### Physiological experiments

Chlorophyll fluorescence measurements were performed by a pulse amplitude modulated fluorometer (Imaging-PAM M-series, Walz, Germany). Experimental conditions for PAM measurements are available in Supplemental Table S5.

Water loss of detached rosettes has been assessed according to Perreau et al., 2020. Hypocotyl elongation assay has been performed according to Végh et al., 2017, and root hair density measurements were accomplished as described by Villaécija-Aguilar et al., 2019 and Meng et al., 2021. The rosette area of 4-week-old plants was measured with ImageJ (Rueden *et al*., 2017). Salt stress treatments were performed according to Lin et al., 2007, and germination percentages have been assayed in 7-day-old seedlings.

### *In vitro* feeding assays with D27 and D27-LIKE1 recombinant proteins

To generate MBP-fused D27-LIKE1 and D27, cDNA inserts without a 37 aa long predicted chloroplast transit peptide (cTPs) were ligated into a BamHI/HindIII-digested pET32b-MBPthr vector (Stráner *et al*., 2018). Plasmids were transformed into *E. coli* BL21(DE3). Starter cultures in 50 ml LB media (100 μgL^−1^ ampicillin) were grown overnight (37*°*C; 180 RPM), then, 1000 ml TB growth culture (supplemented with 100 μgL^−1^ ampicillin and 4 g glucose) was inoculated with 10 ml of starter culture. The cells were grown (37*°*C; 180 RPM) to OD600 of 0.6. To induce protein expression, 0.4 mM IPTG was added. At the same time, the cultures were also supplemented with cysteine (2 mM) and a mixture of FeSO_4_, FeC_6_H_5_O_7_, and C_6_H_8_O_7_xFe_x_NH_3_ (all purchased from Merck KGaA, Darmstadt, Germany) 0.2 mg mL^−1^ each (Jaganaman *et al*., 2007) to ensure high yield of active recombinant Fe-S protein. After 12 hrs of incubation at 18 *°*C, cells were harvested by centrifugation (4500 g, 30 min, 4°C). Cells were resuspended in buffer “A” (50 mM HEPES, 150 mM NaCl, 5 mM TCEP, pH 7.4) and lysed by sonication. The lysate was clarified by centrifugation (23000 g, 30 min, 4°C). The supernatant was applied to a 5 ml MBPTrap HP Column (Cytiva, Marlborough, MA, USA) using an FPLC system (Bio-Rad NGC), pre-equilibrated and washed with 10 CV of buffer “A”. The MBP-fused proteins were eluted with buffer “A” supplemented with 40 mM maltose and were dialyzed immediately against 5L buffer “A” overnight at 4°C. The purity of the collected fractions were checked by SDS-PAGE (Supplementary Figure S4.). All feeding experiments were carried out in four biological replicates.

*In vitro* assays were performed in a total volume of 200 µl. 50 µg enzyme (in 50 µl) was incubated with 40 µM substrate (evaporated to dryness under N_2_ atmosphere and dissolved in 50 µl MQ after micelle-solubilization in EtOH containing 0.8% TritonX-100) in 100 µl incubation buffer containing 1 mM TCEP, 100 mM HEPES (pH 7.4), 0.1 mM FeSO_4_ and 1 mg mL^-1^ catalase in dark for 1 hour at 28*°*C, 100 RPM.

### Measurement of plant pigments, carotenoids from *in vitro assays* and ABA by UPLC-PDA-MS/MS or HPLC

Determination of ABA, plant pigments and carotenoids were executed on a Waters Acquity I-class UPLC system (Waters Co., Milford, MA, USA) with PDA module coupled to a Xevo TQ-XS tandem mass spectrometric system equipped with a UniSpray™ source. Also, pigment determination has been accomplished by HPLC (Shimadzu Prominence; Kyoto, Japan). Sample preparation, as well as UPLC and HPLC method details and the respective MRM transitions used for quantification are available in Supplementary Table S5-S6.

### Statistical Analysis

ANOVA and *post hoc* comparisons of means were performed with OriginPro software (OriginLab Corporation, Northampton, MA, USA) using Tukey’s HSD test.

## Acknowledgements

The authors gratefully acknowledge funding support from the Hungarian National Research, Development and Innovation Office (NKFIH Grants OTKA FK128637, K128644 and GINOP-2.3.2-15-2016-00029). This work was completed within the framework of the ELTE Thematic Excellence Program 2019 by the NKFIH under project “Szint+”. We thank Attila Fábián (AI-CAR) for confocal imaging, Mihály Dernovics (AI-CAR) for carotenoid isomerisation and Mark T. Waters (University of Western Australia) for the helpful comments on the manuscript.

## Author contributions

ZG, BM, IN, EB, ED and VS performed physiological experiments; ZG, BM and SV created constructs and performed transformations; NI and BM constructed phylogenetic tree; ZG, BM and VS performed in vitro experiments, KÁH, VN and LK extracted carotenoids and run UPLC/HPLC experiments; PS and AP purified recombinant proteins. ZG and VS conceived the study and provided supervision. ZG, BM and VS wrote the manuscript with contributions from all authors.

## Supplemental Data

**Supplementary Figure S1**. Consensus sequences of the phylogenetic tree.

**Supplementary Figure S2**. Sequence logo of the highly conserved D27 domain (DUF4033) in all four D27-like proteins.

**Supplementary Figure S3**. Abruption of D27-LIKE1 does not affect the function of the photosynthetic apparatus.

**Supplementary Figure S4**. A, SDS-PAGE electrophoresis of MBP-fused D27-LIKE1 and D27 recombinant proteins. B, Pelleted recombinant D27-LIKE1 and D27 protein display brown colour.

**Supplementary Figure S5**. Chromatograms of feeding experiments.

**Supplementary Figure S6**. ABA and PA content in 2-week-old detached rosettes of *d27-like1-1* and wild-type after 20% water loss.

**Supplementary Figure S7**. qRT-PCR analysis of *D27-LIKE1* in overexpressing lines.

**Supplementary Table S1**. Extracted D27-like sequences used for phylogenetic analysis.

**Supplementary Table S2**. Carotenoid content of 4-week-old detached rosettes of *d27-like1-1*, overexpressing and complemented lines.

**Supplementary Table S3**. Enzymatic activity of recombinant D27-LIKE1 and D27 protein *in vitro*.

**Supplementary Table S4**. List of primer sequences used in this study. **Supplementary Table S5**. Sample preparation and detailed UPLC and HPLC methods. **Supplementary Table S5**. UPLC methods and MRM transitions.

