## Supplementary Figure for "D27-LIKE1 carotenoid isomerase has a preference towards *trans*/*cis* and *cis*/*cis* conversions in Arabidopsis"

### Supplementary Figures

#### Supplementary Figure S2.

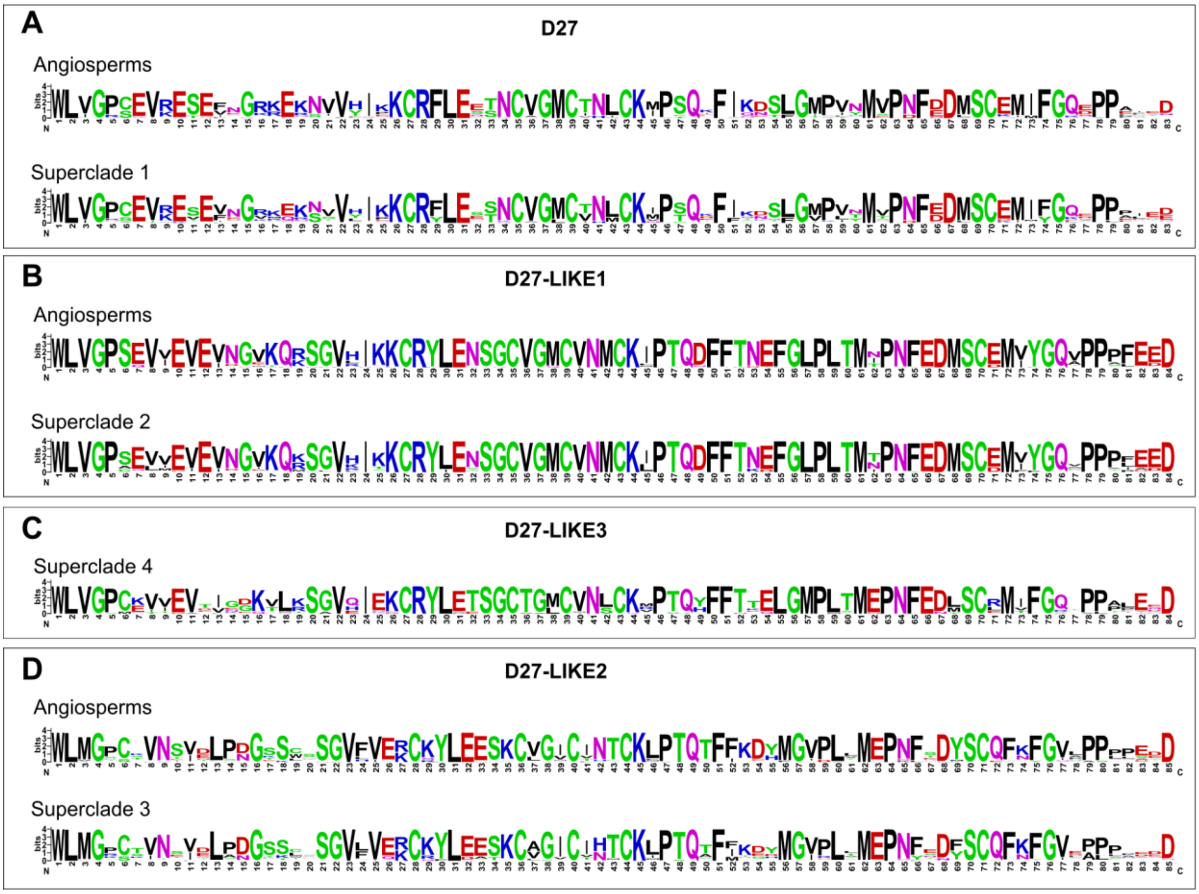

**Supplementary Figure S3.**

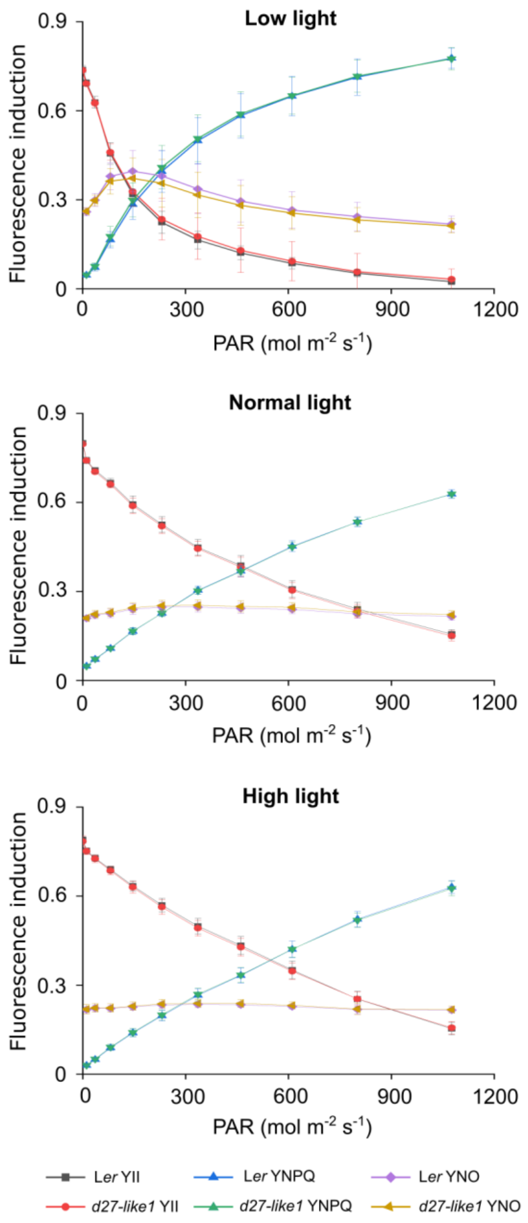


**Supplementary Figure 2.** Abruption of *D27-LIKE1* does not affect the function of the photosynthetic apparatus. Chlorophyll fluorescence analysis of *d27-like1-1* has been assessed by measuring Y(II), Y(NPQ) and Y(NO) parameters of 4-week-old L*er* and *d27-like1* genotypes after growing one week under low (10 μmol $m^{-2}$ $s^{-1}$), normal (100 μmol $m^{-2}$ $s^{-1}$) and high light (800 μmol $m^{-2}$ $s^{-1}$) regimes. Each value represents the mean ± SD of 5 replicates.

### Supplementary Figure S4.

Supplementary Figure 4.

SDS-PAGE electrophoresis of MBP fused proteins (D27-LIKE1 and D27)

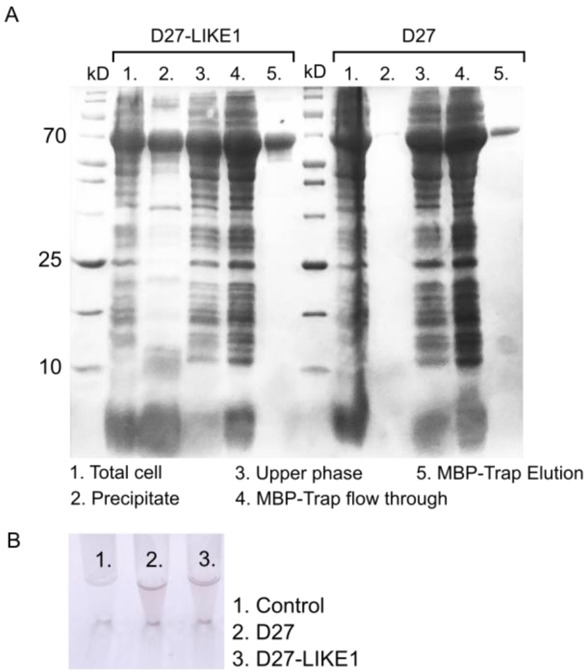

**Supplementary Figure S6.**

Supplementary Figure 6. ABA and PA content in 2 -week-old detached rosettes of d27-like1 and wild-type after 20% water loss.

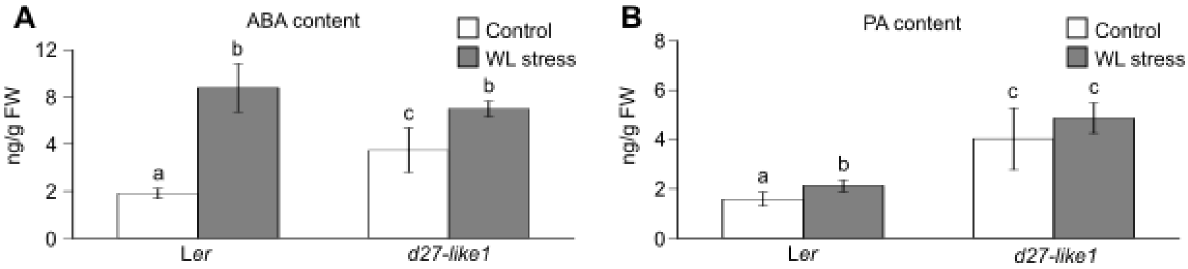

**Supplementary Figure S7.**

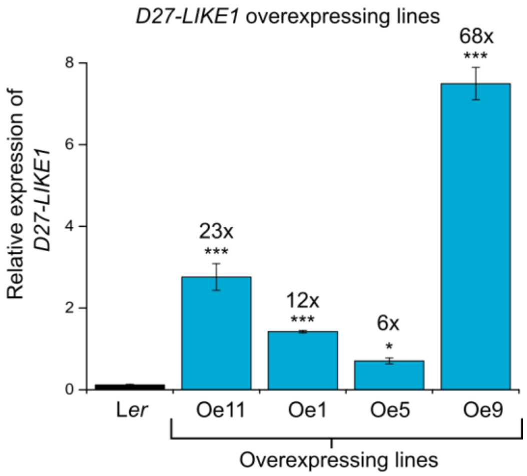
